## Supplemental Figures for "Ancient multiplicity in cyclic nucleotide-gated (CNG) cation channel repertoire was reduced in the ancestor of Olfactores before re-expansion by whole genome duplications in vertebrates"

### CNGA neighbours

Figures S1-S7

### S1: ENSFM00250000000562 - ELF

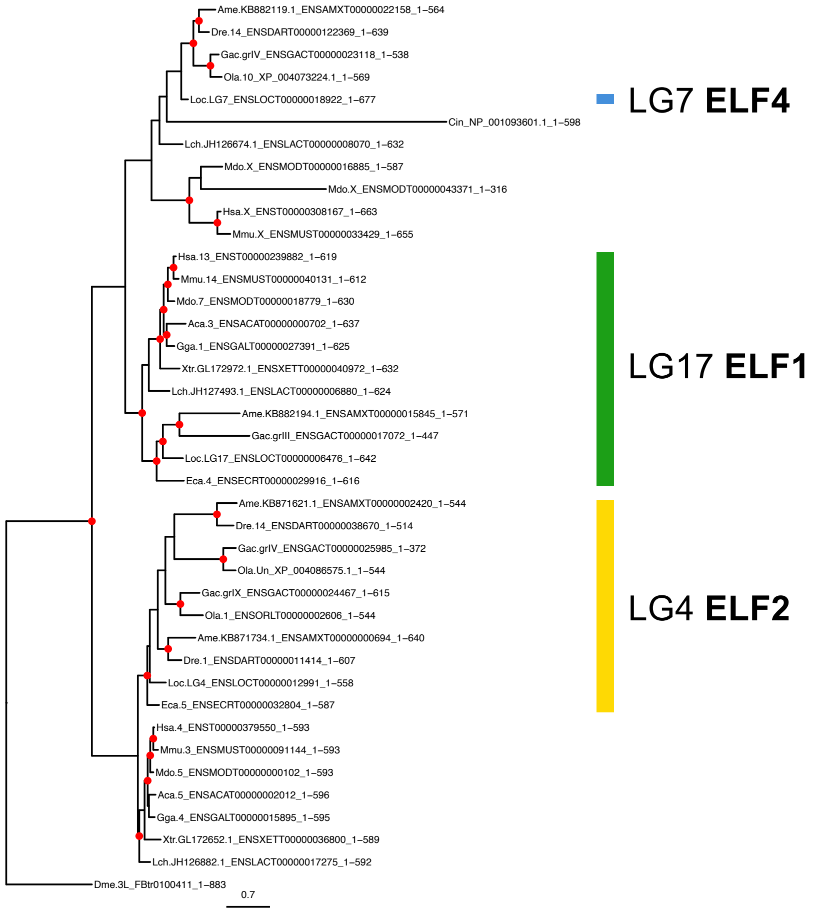

### S2: ENSFM00260000050545 - KCTD

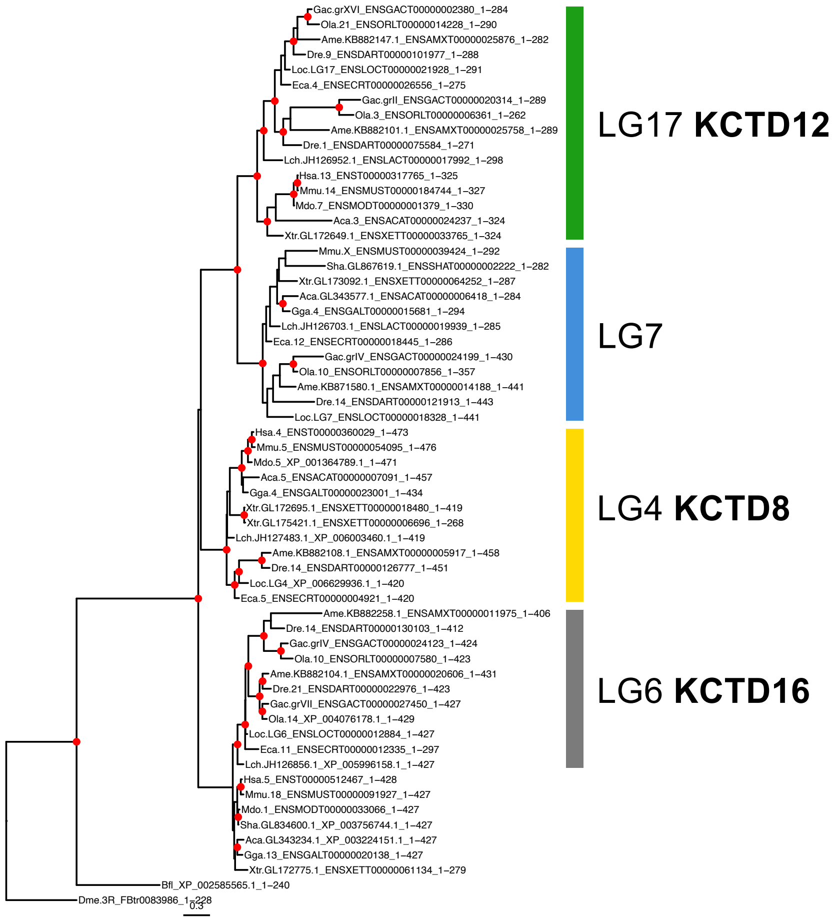

S3: ENSFM00280000058686 - BMX

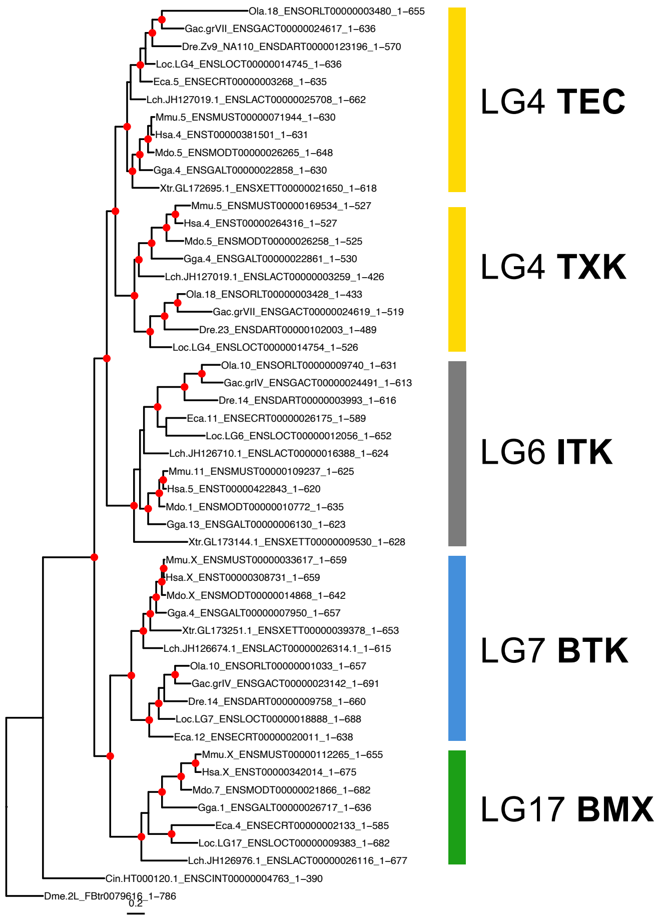

### S4: ENSFM00500000269879 - EDNR

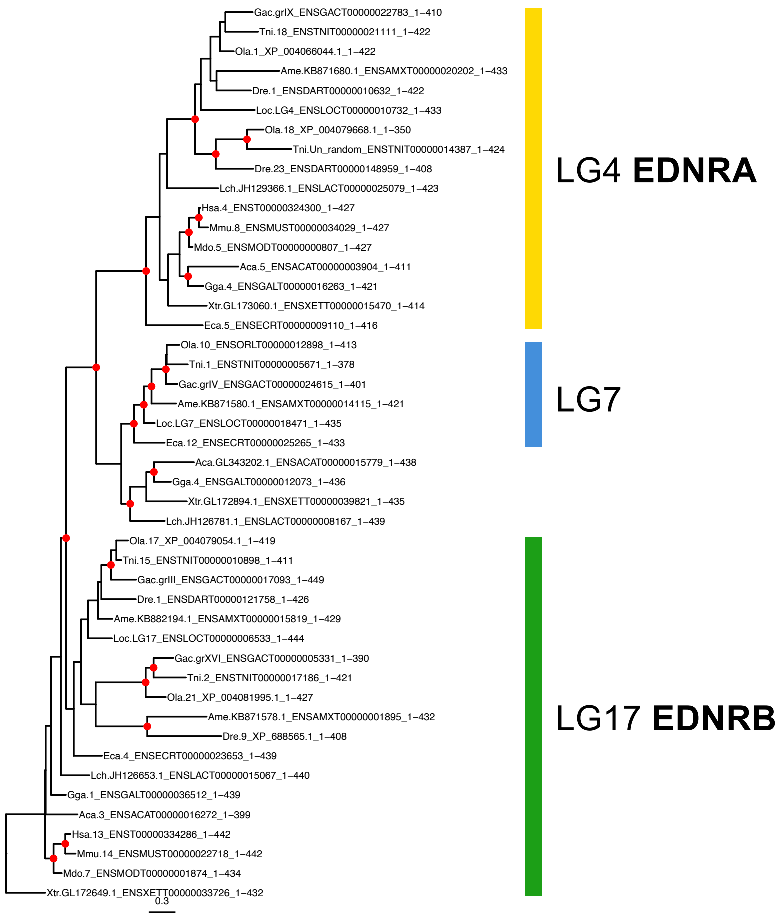

### S5: ENSFM00500000270315 – RAB33

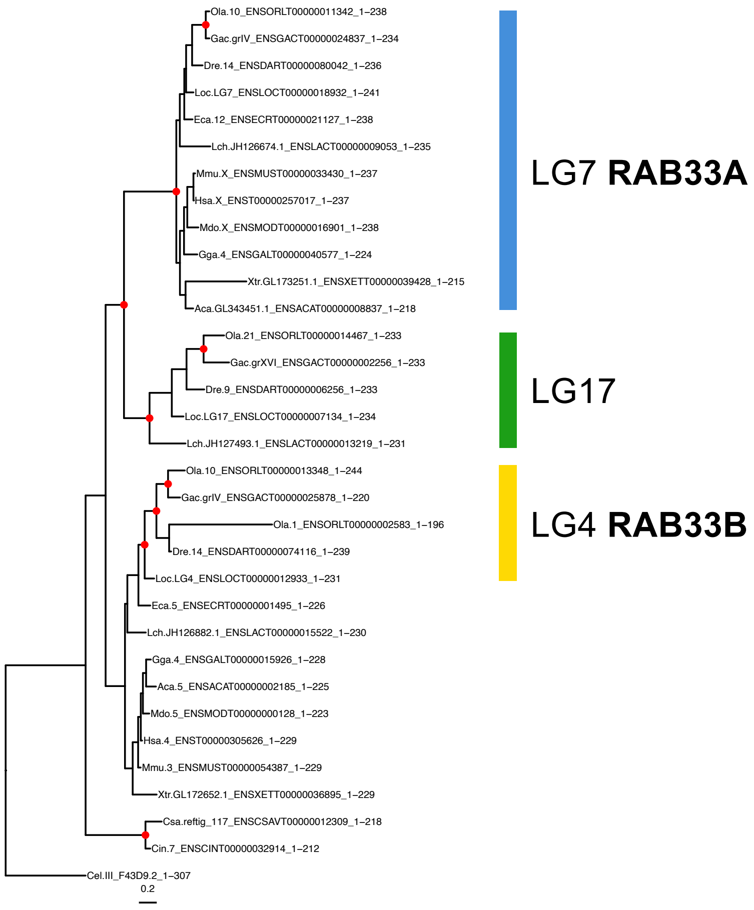

### S6: ENSFM00730001521426 - PCDH

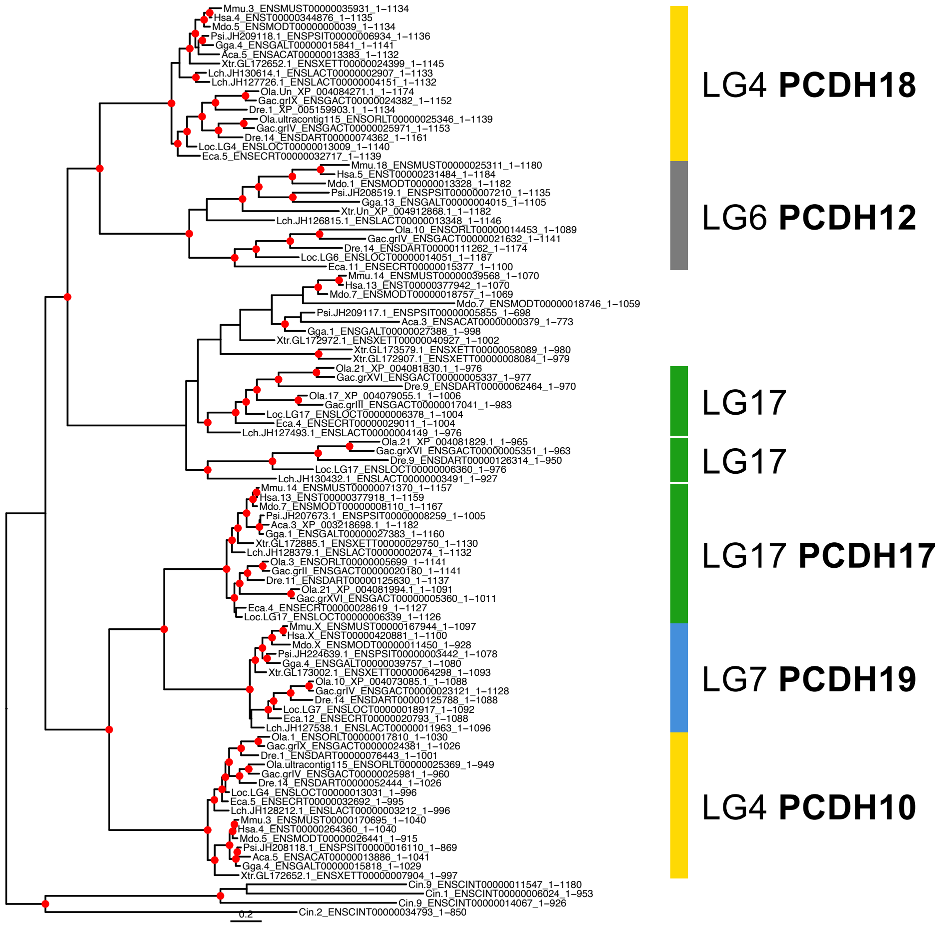

### S7: ENSFM00730001521603 - NIPA

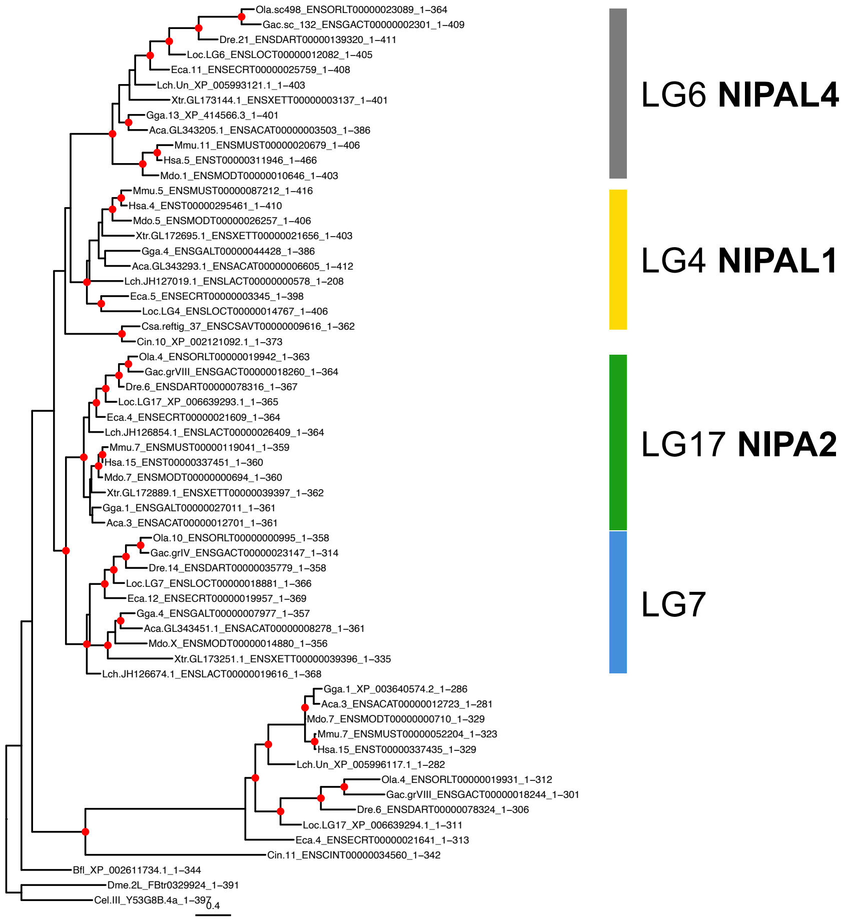

### CNGB neighbours

Figures S8-S17

### S8: ENSFM00250000000331 – SLC7A

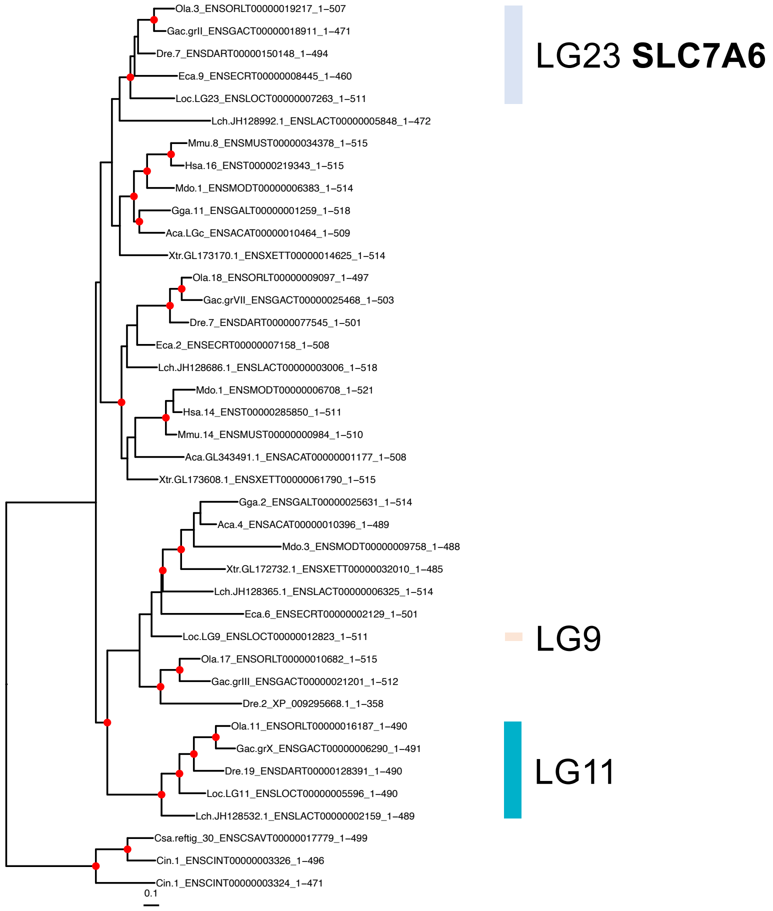

### S9: ENSFM00250000001569 - RANBP

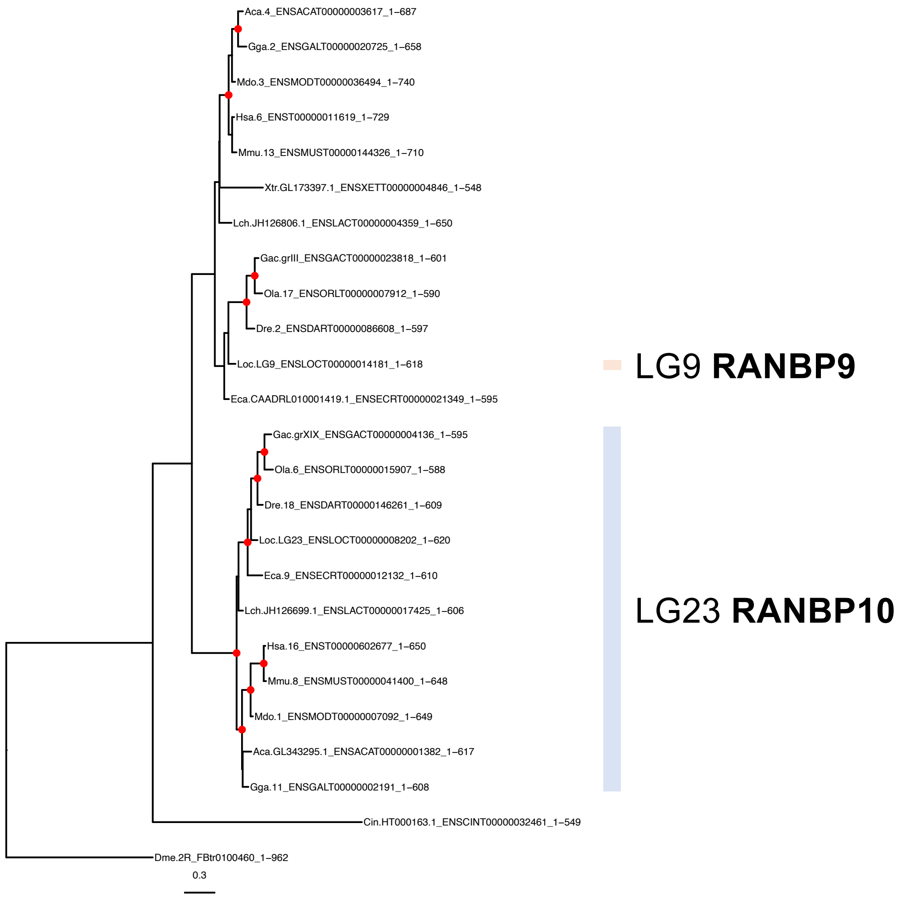

### S10: ENSFM00250000001904 - PHLPP

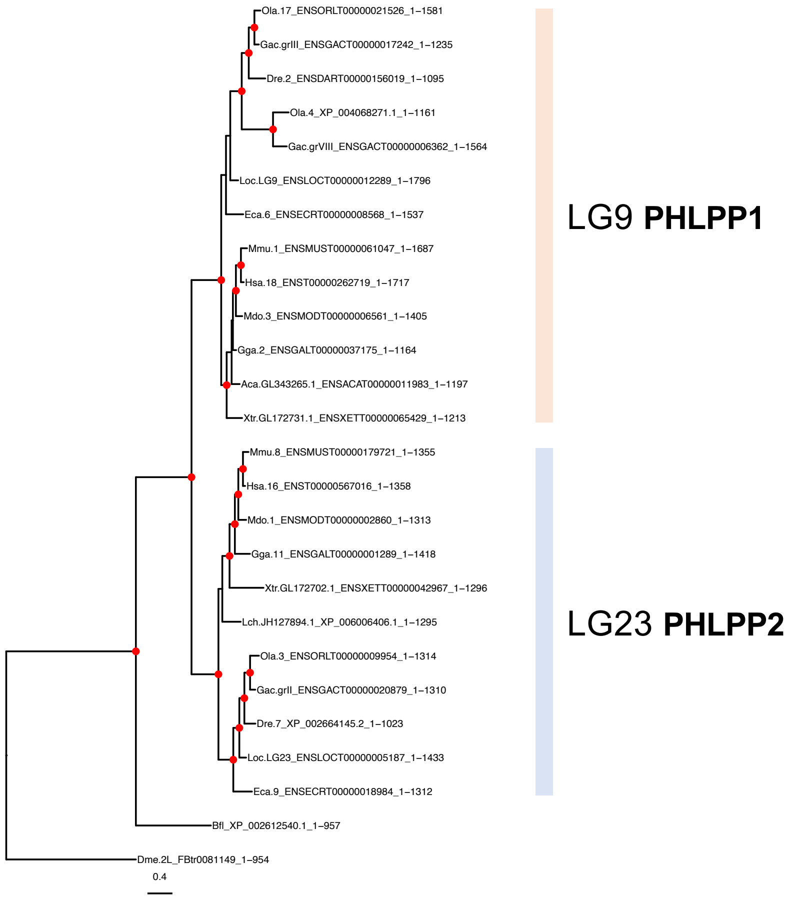

### S11: ENSFM00250000002105 - GPT

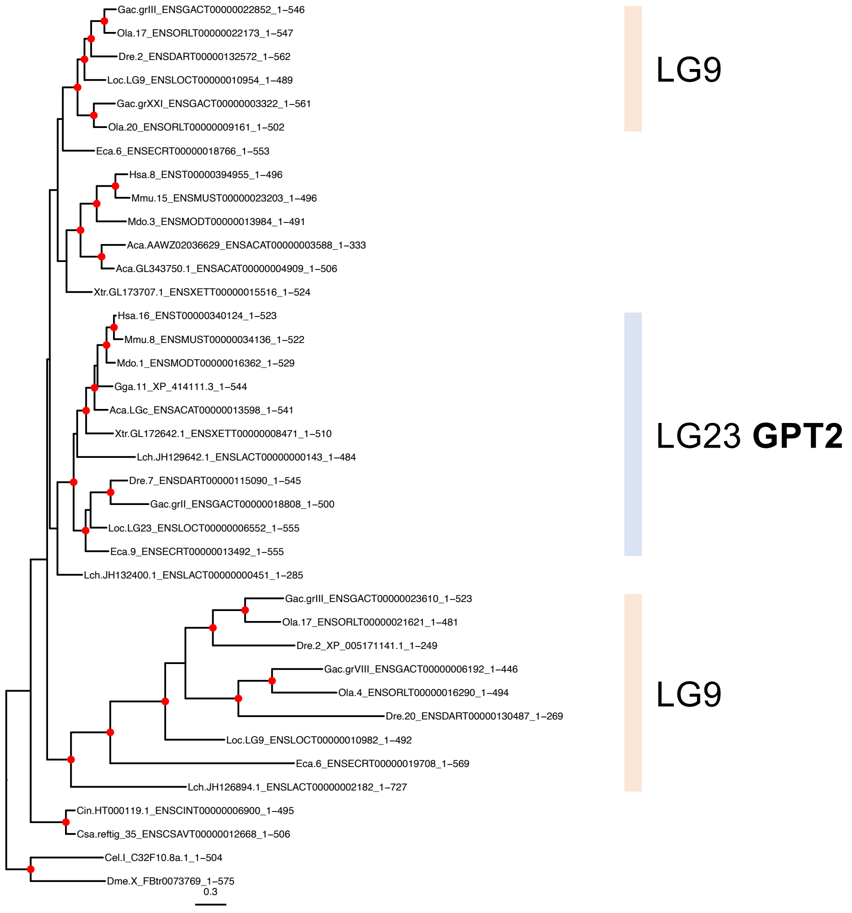

### S12: ENSFM00250000003242 - ATP6V0D

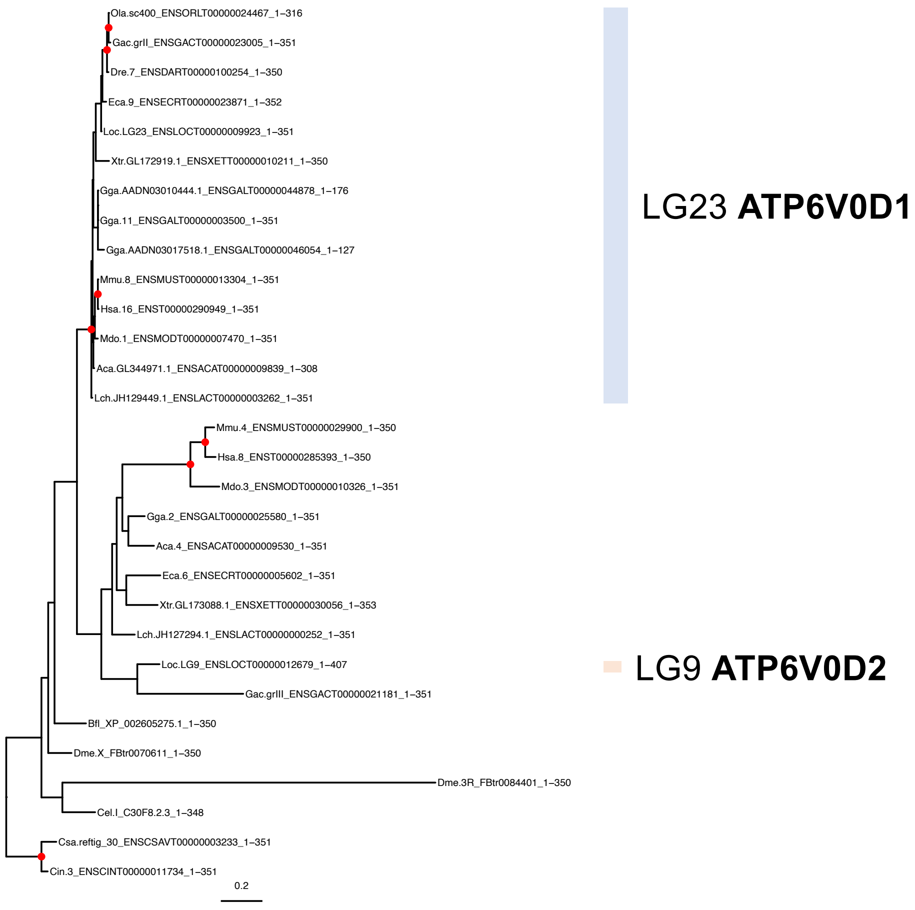

### S13: ENSFM00250000003657 - GFOD

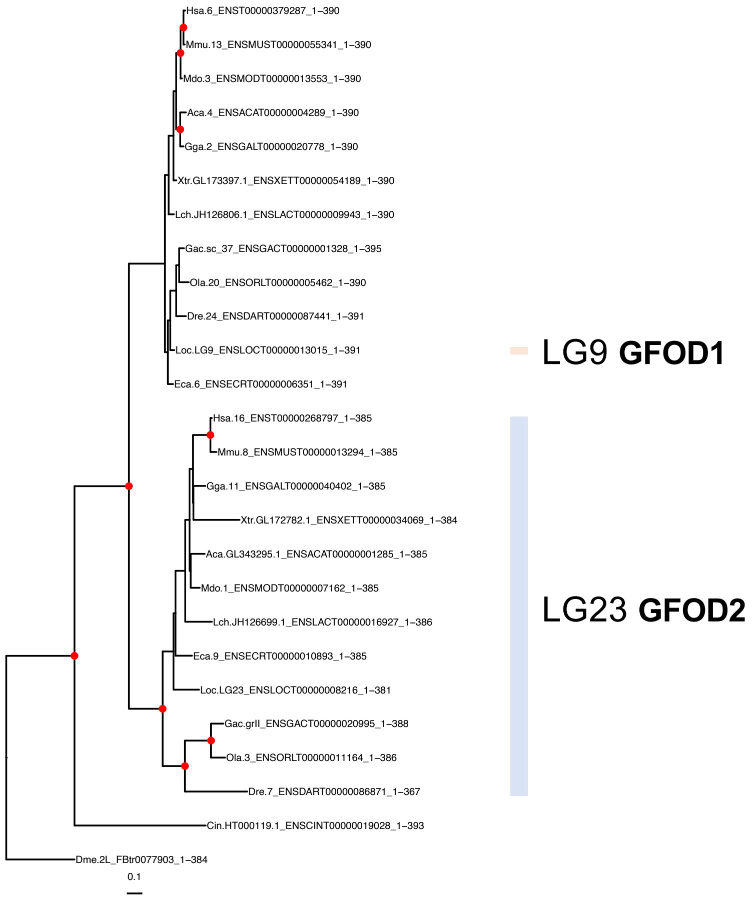

### S14: ENSFM00260000050376 – SLC12A

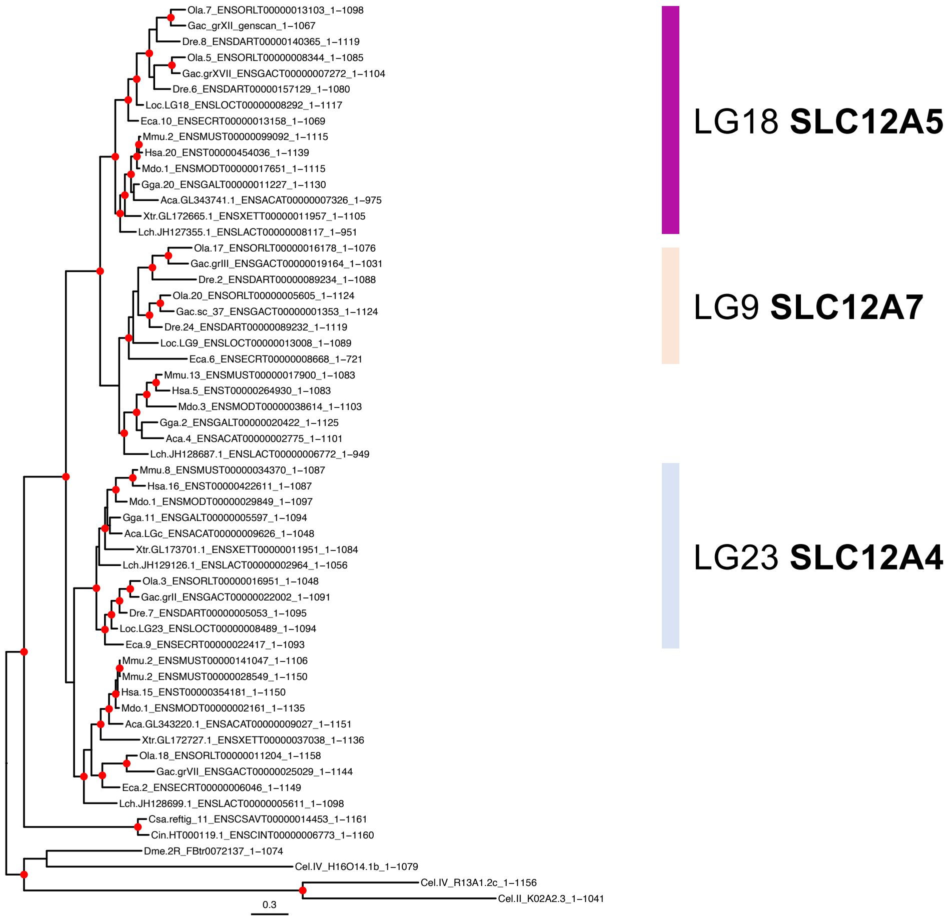

### S15: ENSFM00400000131720 - MMP

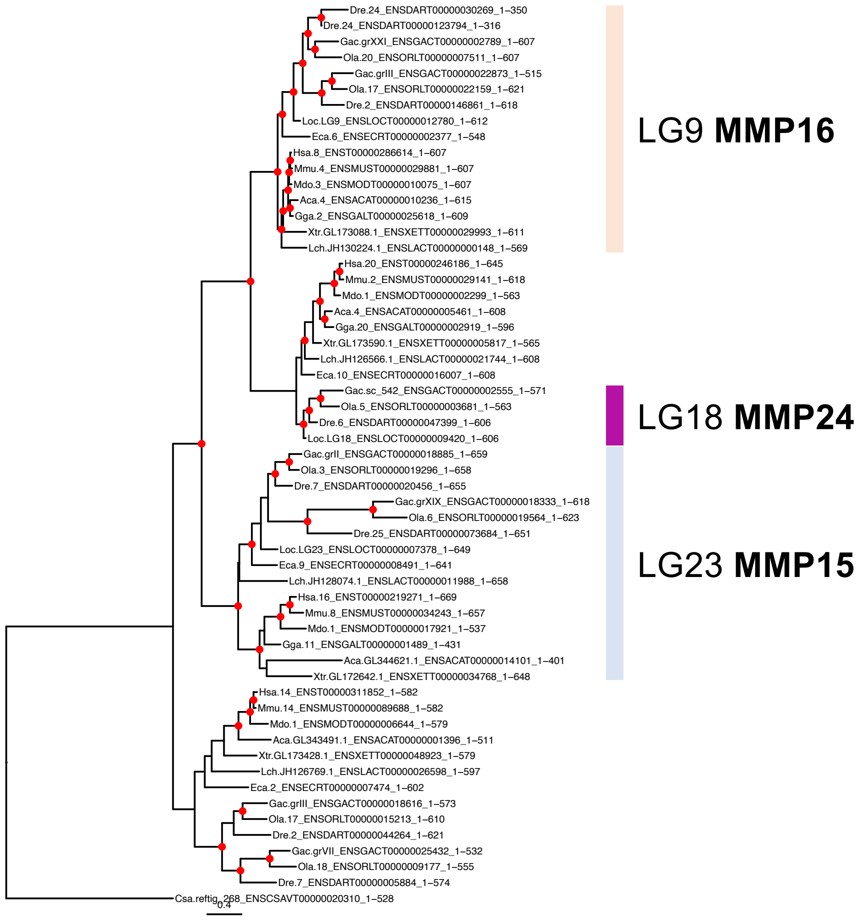

S16: ENSFM00730001521337 - CPNE

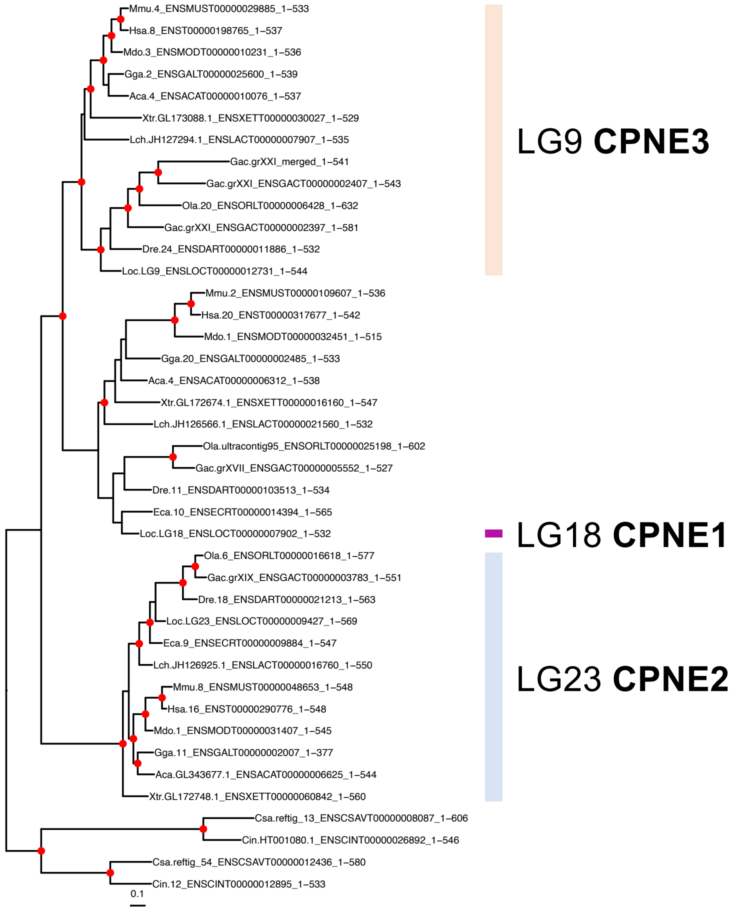

### S17: ENSFM00730001521655 - WWP

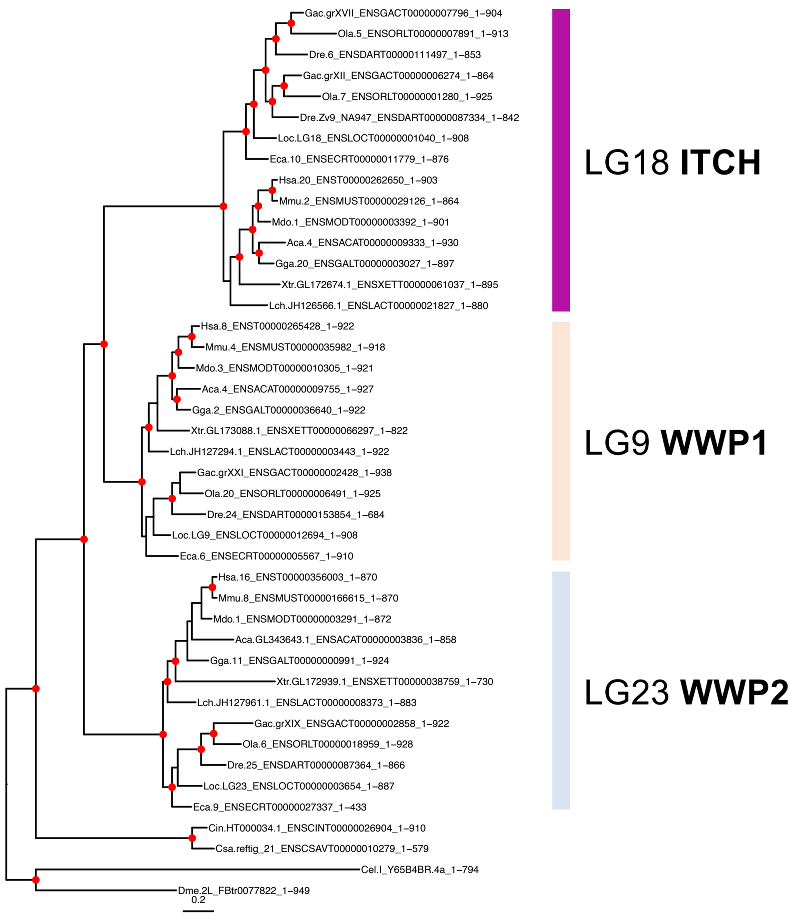

S18:

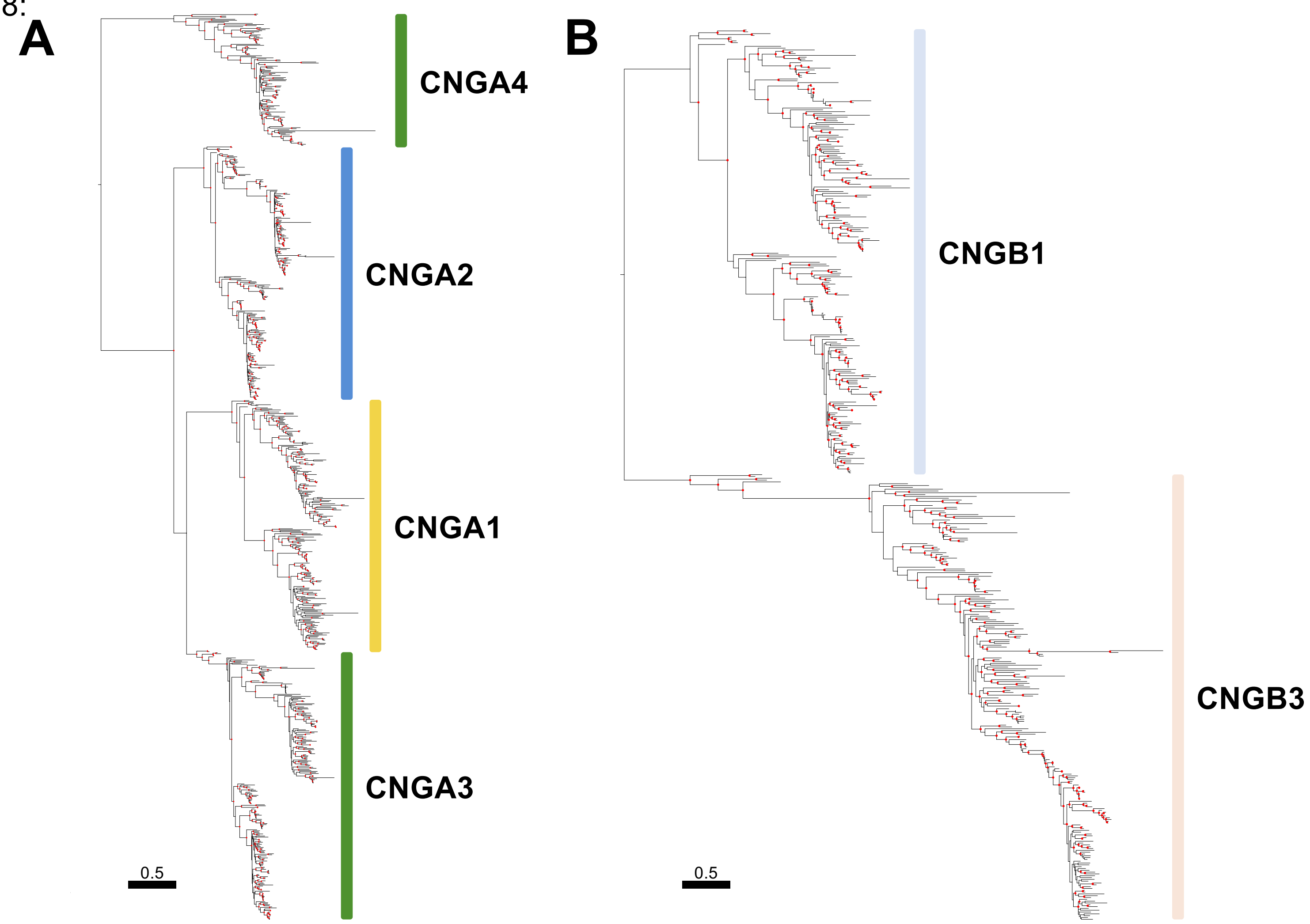

S19:

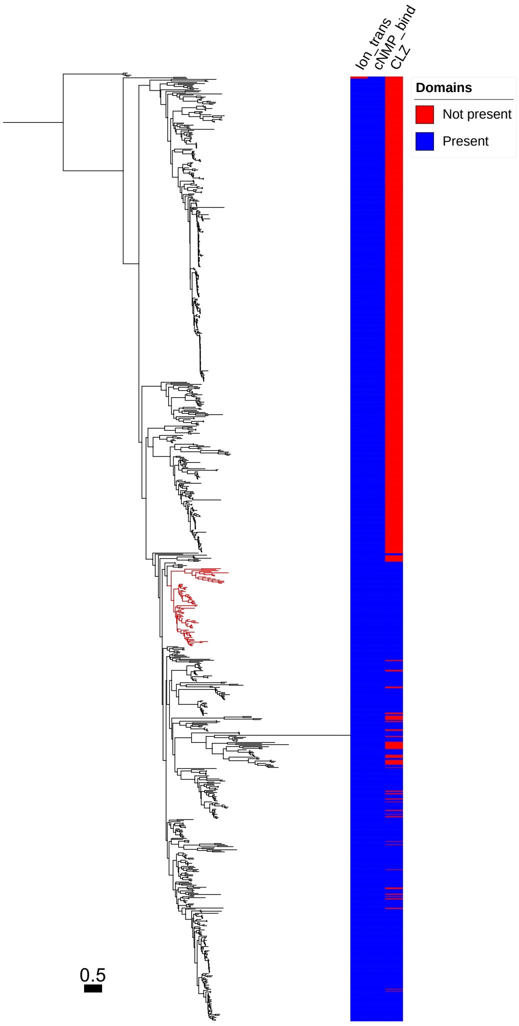
